# A genome-scale metabolic model of a pathosystem sheds new light on bacterial wilt

**DOI:** 10.1101/2024.12.12.628148

**Authors:** Léo Gerlin, Stéphane Genin, Caroline Baroukh

**Affiliations:** Univ Toulouse, INRAE, CNRS, LIPME, Castanet-Tolosan, France; INRAE, INSA Lyon, BF2I, UMR203, 69621 Villeurbanne, France

**Keywords:** Genome-scale metabolic model, plant-pathogen interaction, tomato, *Ralstonia solanacearum*, putrescine

## Abstract

During plant infection, complex metabolic interactions occurs between host and pathogen, including a genuine competition for resources. While the pathogen exploits host nutrients to support its growth and virulence, the plant attempts to restrict pathogen multiplication by limiting nutrient availability or producing antimicrobial compounds.

To unravel these trophic interactions, we constructed a genome-scale metabolic model of a complete pathosystem by integrating a multi-organ metabolic model of the plant, a pathogen metabolic model, quantitative measurements, and a mathematical framework based on sequential flux balance analyses (FBAs). This strategy was applied to the *Ralstonia pseudosolanacearum*-tomato system.

For the first time, quantitative fluxes of matter occurring during a plant infection were predicted. The model shows that (i) plant photosynthetic capacity is a stronger constraint than mineral availability for bacterial proliferation, (ii) infection-induced reduction of plant transpiration limits and ultimately halts first plant growth, then pathogen expansion, (iii) stem resource hijacking can enhance bacterial growth but remains secondary, and (iv) pathogen-excreted putrescine is likely reused for the plant’s needs.

This study delivers the first holistic and quantitative representation of trophic interactions within a plant-pathogen system and highlights the central importance of water flow when the infectious agent is a fast-growing, xylem-colonizing bacterium.

## Introduction

Metabolism is a crucial parameter at the center of interactions between plants and pathogens. Indeed, during host infection, pathogens must efficiently find resources in hostile (and sometimes nutrient-poor) environments to sustain both their growth and virulence (Abu Kwaik & Bumann, 2013). This implies a trade-off for the infecting pathogen, which must balance its carbon and energy expenditure between two objectives: on one hand, limiting plant defense and hijacking plant resources, and on the other hand, supporting its own multiplication and proliferation (Berger et al., 2007). On the plant side, a similar trade-off exists where the plant must distribute resources between maintenance/growth (to ensure reproduction) and immunity, while coping with an increasing infecting population that affects its physiology (Herms & Mattson, 1992; Walters & Heil, 2007). Understanding the complex metabolic interactions occurring between the plant and the pathogen is essential for gaining a holistic view of the infection process. In particular, this knowledge could help to clarify (i) what drives the altered plant phenotype during infection (e.g., growth arrest), (ii) which environmental factors (light, minerals, etc.) limit pathogen growth and population density, (iii) whether the pathogen is able to reprogram the host to allocate nutrients for its own benefit, as *Agrobacterium tumefaciens* does with opines (Zupan et al., 2000), and (iv) how plant metabolic pathways are affected by the presence of the pathogen, especially if the pathogen excretes metabolites. However, the use of a simple metabolomic comparison between infected and non-infected plants is not enough to answer these biological questions, in particular because pathogen metabolism is nearly indistinguishable from plant metabolism, except for very specific metabolites, and metabolism results from the dynamic interplay of multiple synthesis and degradation processes. To better tackle this interaction, a systemic and computational approach is necessary. Ideally, this approach should be quantitative, since only by assigning numerical values can one capture orders of magnitude, thresholds, and trade-offs that would otherwise remain invisible in purely qualitative reasoning.

To this end, we propose in this study to use genome-scale metabolic models (GSMM). GSMMs integrate the metabolic potential of an organism of interest, encompassing thousands of metabolic reactions based on genomics, physiological and biochemical evidences. These models generate quantitative estimations of how matter flows through the biological system under investigation (Gu et al., 2019) as well as phenotypic parameters such as growth. GSMMs have repeatedly proven to be pertinent tools for unraveling complex metabolism, making them suitable for elucidating plant - pathogen metabolic interactions (Gerlin, Frainay, et al., 2021).

Recently, genome-scale metabolic modeling has been applied to the plant – pathogen system of tomato and *Phytophthora infestans* (Rodenburg et al., 2019). It allowed to identify potential pathogen’s targets and plant pathways to engineer in order to block the infection process (Gerlin, Frainay, et al., 2021) but the model developed only provided qualitative insights by exploiting connectivity and topology. It also did not predict the effect of the presence of the pathogen on plant physiology.

A quantitative model of the plant – mutualist symbiont system *Medicago truncatula* - *Sinorhizobium meliloti* has also been recently developed (Pfau et al., 2018; diCenzo et al., 2020). Yet modeling a pathogenic interaction, as compared to a mutualistic one, raises additional methodological questions: i) the choice of the objective function (i.e. the flux(es) to minimize/maximize) in an antagonistic relationship, given that the modeling approach assumes an optimal behavior of the biological system; ii) the need for extensive data acquisition, including the plant/pathogen ratio, growth rates, and metabolites available for the pathogen. These considerations are crucial for developing accurate, meaningful and quantitative models of plant-pathogen interactions.

In this study, we constructed the first quantitative genome-scale metabolic model of a plant–pathogen interaction. For our model pathosystem, we used *Ralstonia pseudosolanacearum* GMI1000 and its natural host, the tomato plant. *R. pseudosolanacearum* is a soil-borne beta-Proteobacterium that enters plants through roots to reach the vascular tissues. The pathogen is then able to reach aerial parts, and proliferate dramatically in xylem vessels, which are typically considered nutrient-poor environments (De La Fuente et al., 2022). Bacterial density has been identified as the cause of xylem vessel clogging (Ingel, Caldwell, Duong, Parkinson, McCulloh, Iyer-Pascuzzi, McElrone, & Lowe-Power, 2022), leading to lethal wilting symptoms. A progressive growth arrest and transpiration decline of the plant has also been observed (Gerlin, Escourrou, et al., 2021). *R. pseudosolanacearum* has been extensively studied for its virulence factors (Vailleau & Genin, 2023) and has gained growing attention for the metabolic aspects of its lifestyle (Peyraud et al., 2016; Lowe-Power et al., 2018; Lowe-Power et al., 2018; Peyraud et al., 2018; Hamilton et al., 2021; Gerlin, Escourrou, et al., 2021). It was among the first plant pathogens having a high-quality GSMM (Peyraud et al., 2016). GSMMs for the tomato plant, *R. pseudosolanacearum*’s natural host, were also developed (Gerlin, Frainay, et al., 2021): a cell genome-scale metabolic model was originally published in 2016 (Yuan et al., 2016) and a genome-scale multi-organ metabolic model, with particular emphasis on xylem vessels (where *R. pseudosolanacearum* is most metabolically active), was recently published (Gerlin et al., 2022). A core metabolic model of tomato stem in the context of a fungal infection has also been developed (Lacrampe et al., 2024).

The quantitative genome-scale metabolic model developed in this study was named VIRION for VIrtual Ralstonia-tomato plant interactION. It was based on the multi-organ metabolic model of the tomato plant, representing different plant organs during vegetative growth (leaf, stem, root with xylem and phloem as exchange compartments), combined with *R. pseudosolanacearum* model represented as an additional compartment connected to the xylem. To accurately simulate the antagonistic interaction between the plant and the pathogen, we employed a set of sequential Flux Balance Analyses (FBAs). To be more accurate, VIRION also integrates a comprehensive set of physiological and metabolic data: i) tomato vegetative growth (organ ratio, growth rates, xylem metabolites), ii) tomato infection by *R. pseudosolanacearum* (plant/pathogen ratios, impacts of infection on physiology), and iii) *R. pseudosolanacearum* physiology and metabolic capacities (growth rates, assimilation and excretion capacities on xylem metabolites).

The model only focused on the trophic aspect of metabolism. It did not account for the plant’s immune response or the pathogen’s virulence, as modeling such phenomena is highly challenging and requires alternative approaches such as dynamic gene regulatory networks or functional association networks (e.g., immunity-related metabolomic measurements) that are not available for our pathosystem. Moreover, the present study addressed an infection stage that, in the case of *Ralstonia*, corresponds to intense multiplication within plant tissues, and thus likely occurs once most of the immune defenses have already been overcome (Gerlin, Escourrou, et al., 2021; Baroukh et al., 2025). At this stage, trophic interactions are therefore expected to be predominant. In addition, this stage of the interaction has received little attention so far, whereas numerous studies have focused on early plant immunity, such as perception and signaling (DeFalco & Zipfel, 2021; Jian et al., 2023).

The model enables to generate quantitative estimation of the growth capacities of *R. pseudosolanacearum in planta* at different stages of the disease, as well as the pathogen’s impact on plant growth and metabolism. We used the model to analyze factors causing the infected plant to cease growth and those limiting bacterial growth. Finally, we also investigated potential hijacking of stem resources during the disease, and the fate of the bacterial excreted putrescine in the plant.

## Results

### Modeling an antagonist plant-pathogen interaction using a multi-compartment metabolic model and sequential Flux Balance Analyses

To model the metabolic interaction between *R. pseudosolanacearum* and the tomato plant, we represented the system as four metabolic compartments corresponding to the main metabolic roles of the plant (leaf, stem, root) and the pathogen. Metabolites can be exchanged between these compartments through xylem and phloem (Figure 1A). Each of these four compartments was represented as a genome-scale metabolic model: iRP1476 for the pathogen (Peyraud et al., 2016) and Sl2183 for the plant compartments (Gerlin et al., 2022) (Figure 1B), with minor modifications as reported in the Methods section. As in Gerlin et al. (2022), the physiological roles of each compartment (described in Figure 1A) were implemented as FBA constraints (Figure 1B). In brief: i) minerals and water can enter the system only through the root and are transported to the upper parts of the plant and to the pathogen exclusively via xylem, ii) photons are assimilated in both the leaf and the stem, with higher efficiency in the leaf, iii) the carboxylase/oxygenase activity of the Rubisco was constrained to experimentally measured values. For a complete description and justification of these constraints, readers are referred to Gerlin et al. (2022).

**Figure 1:**
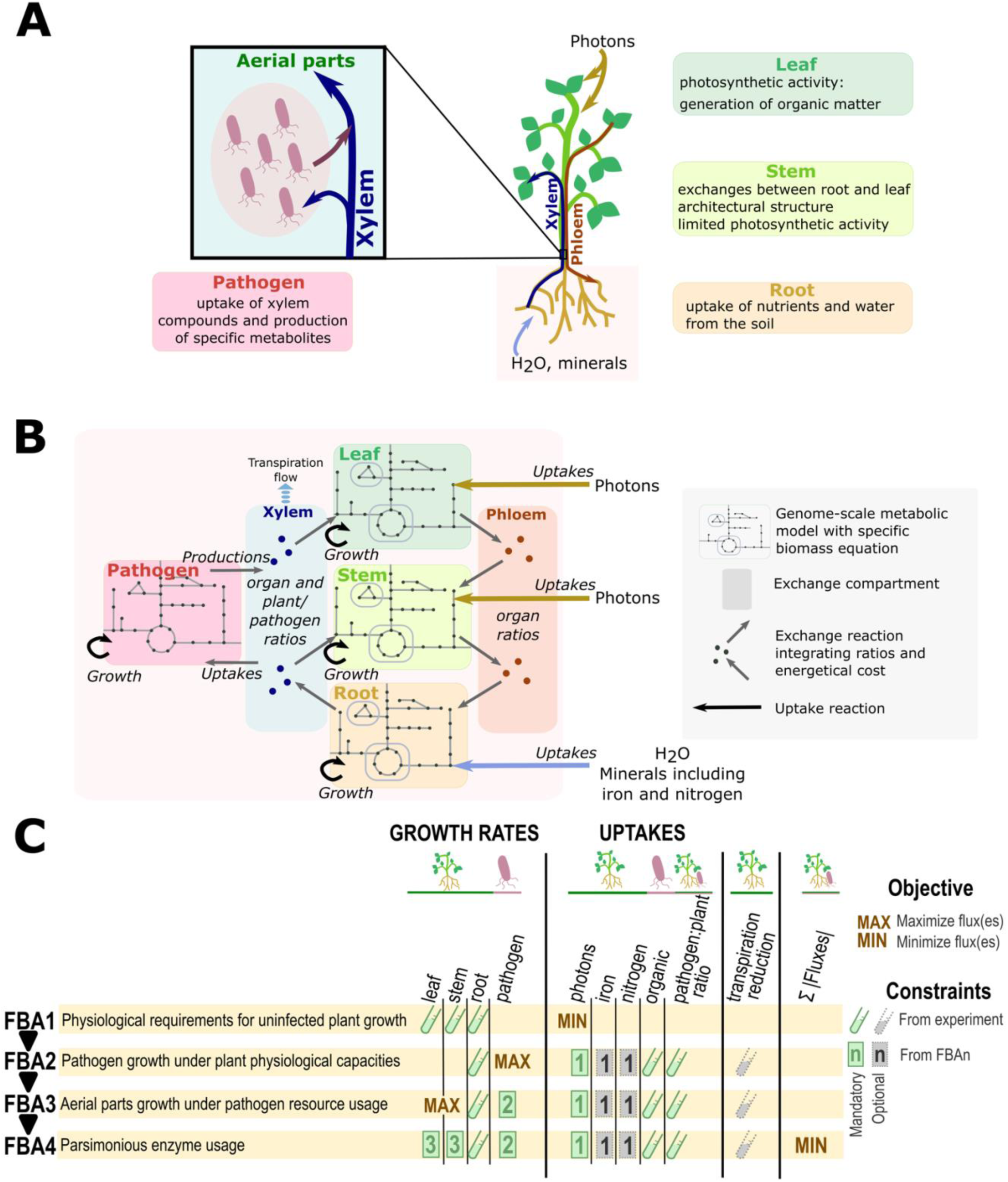
Schematic representation of the metabolic model VIRION. A. The main components of the system modeled in this study. Leaf, stem, root and the pathogen are each represented by a genome-scale metabolic model. Xylem and phloem are exchange compartments. Inputs of the system are photons, connected to leaf and stem, and water and minerals, connected through the root. B. Schematic representation of the multi-compartment model and quantitative approach used in this study. Arrows represent relationships between the different organs and the possible quantitative constraints which can be set in the model (growth, uptakes, exchanges, transpiration flow). C. Sequential Flux Balance Analyses (FBAs). Schematic representation of the different FBAs solved in a sequential approach to obtain a phenotype prediction of the pathosystem (growth rate of plant organs and pathogen) as well as fluxes of matter. The constraints from experiments are detailed in Supplementary Table 1.

FBA is an approach based on an optimization problem (typically formulated as linear programming), in which one or several objective reactions are maximized or minimized to compute metabolic fluxes (Orth et al., 2010). Usually, when modeling ecosystems, the organisms composing the system are assumed to collectively maximize global biomass (diCenzo et al., 2020). However, this assumption is not appropriate for our antagonistic system, in which the pathogen grows at the expense of the plant, leading to decoupled growth dynamics between the two partners. Moreover, in our system, the metabolic composition of xylem sap is severely altered upon bacterial infection, particularly at high bacterial densities (Gerlin, Escourrou, et al., 2021), thereby challenging the quasi-steady state assumption (QSSA). To model this antagonistic relationship, we employed a system of sequential FBAs (Figure 1C). In this approach, an initial FBA determines one or several optimal flux(es), which are subsequently reinjected into the next FBA as additional constraints. A total of four successive FBAs were performed for each bacterial density to predict both bacterial and plant metabolism during the interaction. This four-step FBA workflow was repeated for several bacterial densities representative of those observed *in planta* (Gerlin, Escourrou, et al., 2021). Bacterial density was accounted for by adjusting the stoichiometric coefficients of the exchange reactions between the xylem and the pathogen for each density (see Methods for details).

The sequential 4-FBA approach consisted of the four following steps:

FBA 1: Estimation of the requirements of a healthy plant in terms of photons, nitrogen and other minerals necessary for an optimal growth of the leaf, stem and root.

FBA 2: Estimation of pathogen growth while constraining plant photon and mineral inputs to the levels obtained in FBA 1. This ensured that resource consumption did not exceed that of a healthy plant. Photon constraints were always applied, whereas mineral constraints were optional. In addition, the transpiration flow – known to be significantly reduced after pathogen inoculation (Gerlin, Escourrou, et al., 2021) and influence xylem exchanges - could be constrained at this step (see Methods for details).

FBA 3: Estimation of leaf and stem growth rates given the pathogen growth determined in FBA 2. In both FBA 2 and FBA 3, the root growth rate was fixed to experimentally observed values, since root growth is minimally affected by xylem colonization according to experimental data on infected tomato plants (Gerlin, Escourrou, et al., 2021).

FBA 4: Estimation of metabolic fluxes in the pathosystem under a parsimonious enzyme usage constraint. This final simulation yielded the flux distribution for a given bacterial density.

These 4 FBAs were performed for bacterial densities ranging from 10^6^ to 10^12^ Colony Forming Unit per gram of plant Fresh Weight (CFU.g^-1^ FW), which are bacterial densities typically measured in the plant during an infection cycle (Gerlin, Escourrou, et al., 2021). The simulation results – including plant and pathogen growth rates and the composition of xylem fluxes upon infection - were compared with experimentally observed behaviors to validate the model and gain insights into the pathosystem.

### Identifying metabolic bottlenecks during the plant – pathogen interaction

We used the VIRION model to assess which plant environmental factors - light, nitrogen, iron or water - constrained the plant-pathogen interaction. The analysis considered these factors individually and in combination (Figure 2A). Because infected plants displayed reduced leaf area compared to healthy ones (Gerlin, Escourrou, et al., 2021), light constraint was treated as mandatory: an infected plant cannot absorb more photons than a healthy plant. While other minerals could be studied, our study focused on nitrogen and iron. Nitrogen source affects the concentration of xylem organic metabolites in tomato (Bialczyk et al., 2004), and both nitrogen and iron were suggested to influence the switch to virulence in *R. pseudosolanacearum* (Bhatt & Denny, 2004; Dalsing & Allen, 2014; Dalsing et al., 2015). Light, nitrogen and iron were constrained on their respective uptake rates from FBA 2. The imposed bounds corresponded to the rates computed in FBA 1 (minimization of plant photons for a healthy tomato plant, Figure 1C). Transpiration was also optionally limited by constraining the upper bound of root to xylem transport fluxes, based on a previously established linear relationship between *in planta* bacterial density and transpiration reduction (Gerlin, Escourrou, et al., 2021) (see Methods for further details).

**Figure 2:**
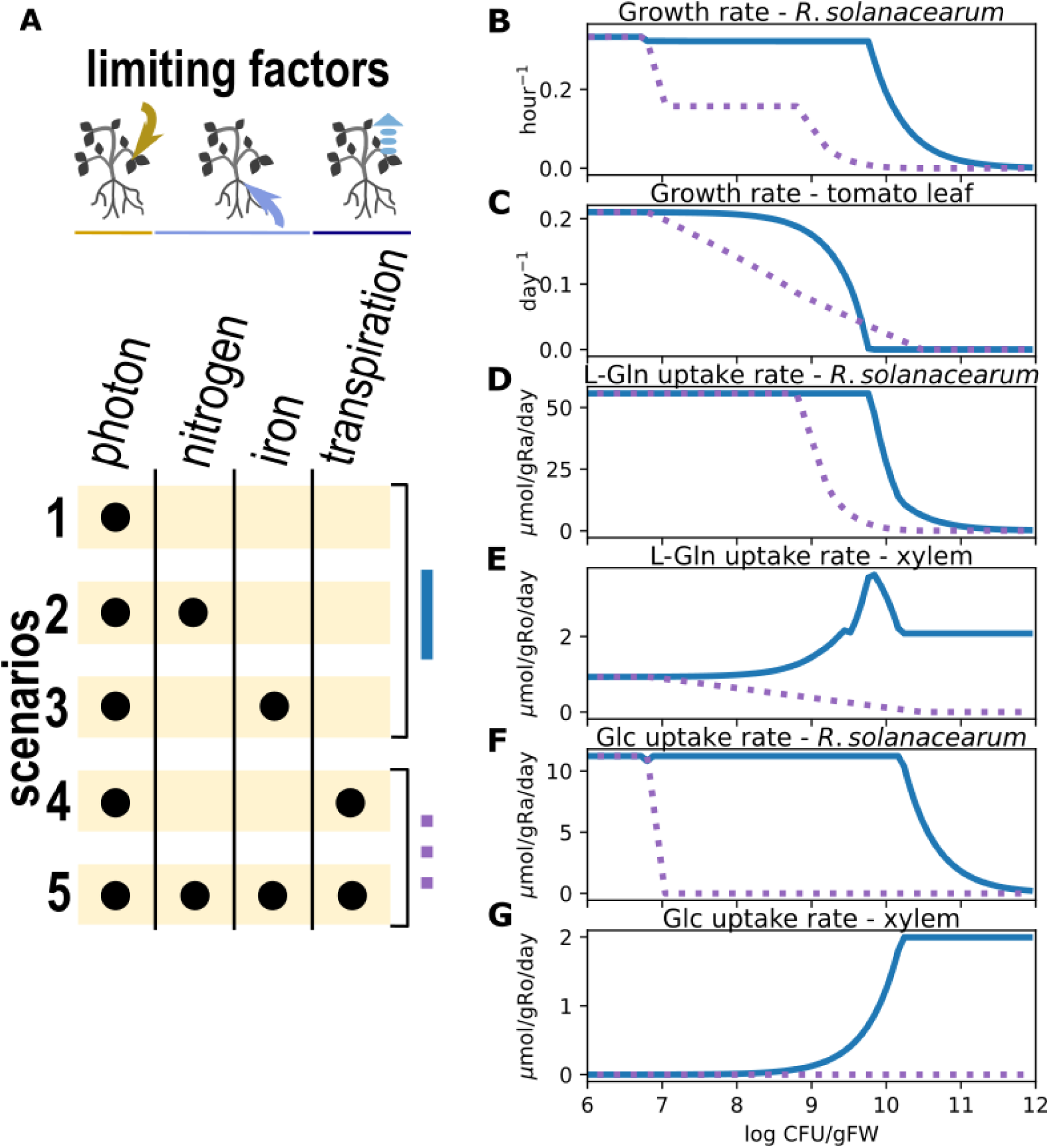
Effect of plant environmental conditions on growth and metabolic exchanges in the pathosystem. A. Simulation scenarios in which different plant environmental conditions were constrained at the FBA 2 step during sequential FBAs. The upper bounds of photons, nitrogen and iron were set to the values obtained from FBA 1. Transpiration constraint was implemented as constraints on the upper bound of fluxes from root to xylem using a linear relationship computed using the experimental values of Gerlin et al. (Gerlin, Escourrou, et al., 2021). B. to G. Effects of plant environmental conditions on growth and metabolic exchanges for bacterial densities ranging from 10^6^ to 10^12^ CFU.g^-1^ FW. gRa: dry weight grams of *R. pseudosolanacearum,* gRo: dry weight grams of root, L-Gln: L-Glutamine, Glc: D-glucose, CFU: colony forming units, FW: fresh weight.

When light is the sole limiting factor (Figure 2), thereby constraining photosynthesis and organic carbon production, *R. pseudosolanacearum* grows at its maximal rate up to a density of 10^9.7^ CFU.g^-1^ FW (Figure 2B). Beyond this threshold, bacterial growth declines sharply, falling below 0.05 h⁻¹ once densities exceed 10^10.6^ CFU.g^-1^ FW. These results indicate that if Ralstonia relies exclusively on biotrophic growth sustained by xylem sap metabolites, the plant cannot support bacterial proliferation beyond roughly 10^10^ CFU.g^-1^ FW. Analysis of the simulated substrate uptake fluxes reveals that glutamine is the primary carbon source sustaining Ralstonia growth (Figure 2D). Glutamine uptake remains high up to approximately 10^10^CFU.g^-1^ FW, beyond which it becomes limiting – first for the growth of the plant’s aerial tissues, and subsequently for Ralstonia. This limitation is temporarily alleviated by glucose uptake, a secondary carbon source that is co-consumed with glutamine. However, at slightly higher bacterial densities (10^10.25^ CFU.g^-1^ FW), the maximal glucose uptake flux in the xylem is also reached, further constraining bacterial growth. These simulation results are consistent with experimental observations: *in planta* analyses and *in vitro* assays have identified glutamine as the principal carbon source of *R. pseudosolanacearum* in tomato xylem sap (Baroukh et al., 2022; Gerlin, Escourrou, et al., 2021; Baroukh et al., 2025). While glucose can moderately support bacterial proliferation, it has been shown to play only an accessory role (Baroukh et al., 2022, 2025; Gerlin, Escourrou, et al., 2021; Hamilton et al., 2021).

However, plant growth capacities were overestimated by the simulations results: optimal leaf and stem growth were maintained in silico up to a bacterial density of 10^8.5^ CFU.g^-1^ FW (Figure 2C), whereas strong growth inhibition was experimentally observed between 10^6^ and 10^8^ CFU.g^-1^ FW (Gerlin, Escourrou, et al., 2021). This discrepancy indicates that additional factors contribute to plant growth arrest. When mineral constraints—specifically iron and nitrogen—were applied, either separately or together, the simulation outcomes were similar to those obtained under light limitation alone (Figure 2). Nitrogen and iron therefore appeared to be less stringent bottlenecks than light. In contrast, limiting transpiration flow produced distinct results, with a decline in both plant and Ralstonia growth rates occurring at lower bacterial densities (Figure 2). Indeed, from approximately 10^7^ CFU.g^-1^ FW, reduced transpiration limited the availability of xylem sap nutrients for both the bacterium and the aerial tissues. Because the model prioritizes Ralstonia growth at the expense of that of the plant, an immediate effect on plant development was observed (Figure 2C).

At higher bacterial densities, xylem fluxes became insufficient to sustain rapid Ralstonia growth, resulting in an intermediate growth rate (Figure 2B) before a complete decline beyond 10^9^ CFU.g^-1^ FW. Interestingly, glutamine remained abundant enough to satisfy the maximal uptake rate of *R. pseudosolanacearum* (Figure 2D), but other metabolites present at low concentrations in the xylem—such as glucose (Figure 2F), sucrose, and amino acids (Supplementary Figure 1)—were no longer assimilated or only at reduced rates, explaining the intermediate growth phase observed. Above 10^9^ CFU.g^-1^ FW, glutamine availability also became limiting, severely impairing bacterial growth and rapidly leading to growth arrest. These findings are consistent with experimental data, which report maximal bacterial densities around 10^9^ CFU.g^-1^ FW (Gerlin, Escourrou, et al., 2021). They also align with predictions from a recently developed macroscopic model of *R. pseudosolanacearum* growth in planta (Baroukh et al., 2025).

Finally, adding nitrogen and iron limitations to the model—already constrained by photon availability and transpiration—did not markedly alter the simulated progression of the disease (Figure 2B–G). This result suggests that the decline in plant transpiration, which reduces xylem fluxes, exerts the strongest influence on both plant and pathogen metabolic capacities. Transpiration decline therefore appears to be the most probable mechanism underlying plant growth arrest, pathogen growth arrest, and the establishment of maximal bacterial density. In contrast, limitations in nitrogen, iron, or carbon (via photon limitation) do not seem to represent the primary drivers of these experimental observations.

### Assessing if hijacking tomato stem resources is a good pathogen strategy

In the previous simulations, the contribution of stem metabolism to xylem sap was forbidden, as no flux from the stem to the xylem was predicted for a healthy plant (Gerlin et al., 2022). However, it can be hypothesized that such fluxes might occur in an infected tomato plant, potentially contributing to pathogen nutrition if the bacterium employs a host resource hijacking strategy. This idea aligns with the hypothesis that certain type III secretion effectors of *R. pseudosolanacearum* can modify plant metabolism (Landry et al., 2020; Macho, 2016).

We re-modeled scenario #5 from the previous section (where photons, nitrogen, iron, transpiration limitations were applied, Figure 2A), this time allowing nutrient export from the stem to the xylem, creating scenario #6 (Figure 3A). Simulation results indicated that permitting the stem to enrich xylem sap had only a slight effect on tomato leaf growth (Figure 3C). In contrast, a more pronounced effect was observed on pathogen growth (Figure 3B): *Ralstonia* could maintain a maximal growth rate up to 10^9.2^ CFU.g^-1^ FW, compared to 10^7^ CFU.g^-1^ FW in scenario #5 (Figure 3B).

**Figure 3:**
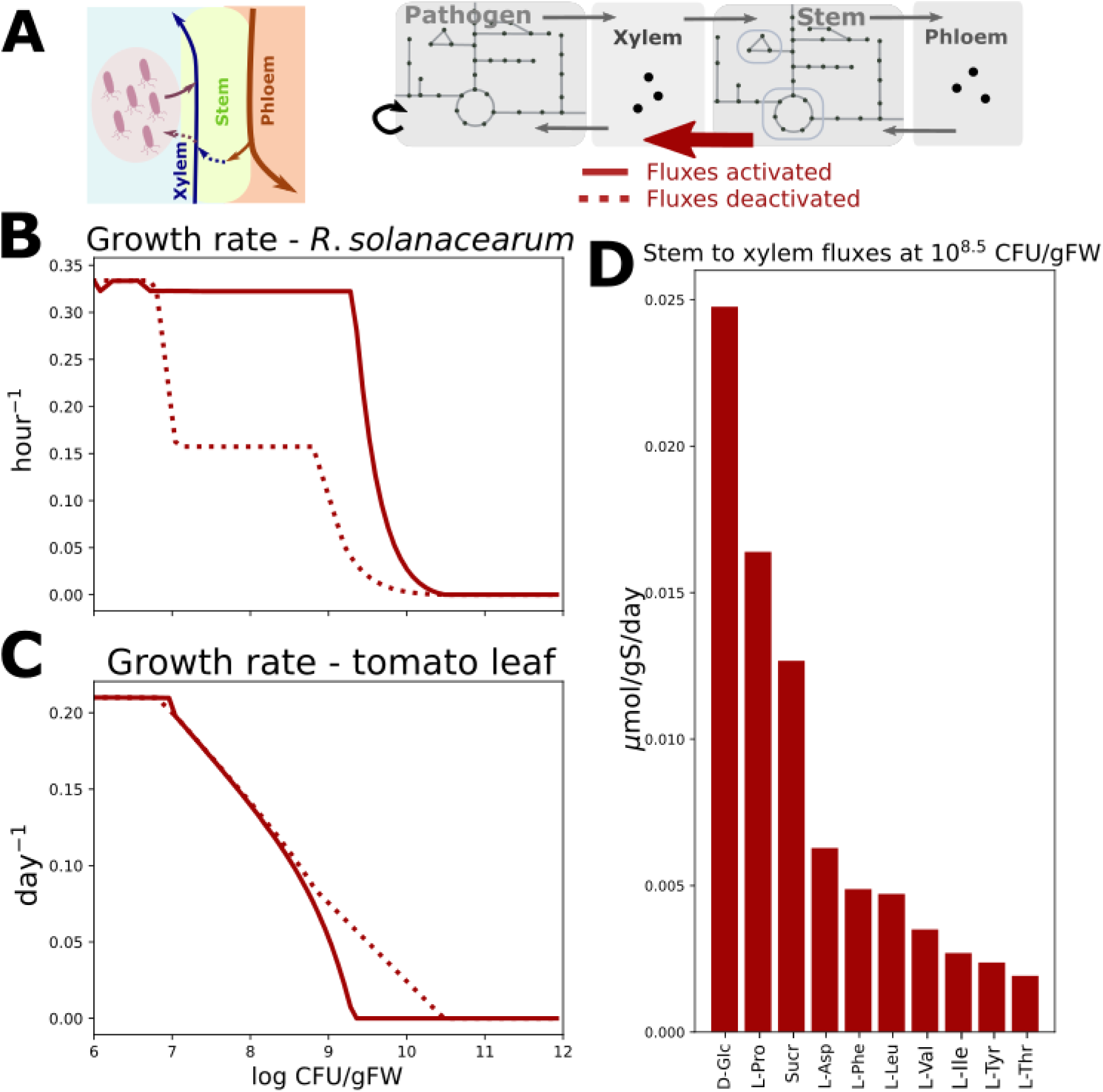
Effect of xylem sap enrichment by the stem on plant growth, pathogen growth and xylem sap composition. A. Schematic representation of the investigated plant fluxes. In scenarios #1 to #5, the stem could not enrich xylem sap. This constraint was lifted to assess its effect on the metabolism of the pathosystem. B. Pathogen growth rate when the stem can (line) or cannot (dotted line) enrich xylem sap. C. Plant growth rate when the stem can (line) or cannot (dotted line) enrich xylem sap. D. Fluxes of matter from stem to xylem when the stem can enrich xylem sap. Fluxes were computed for a bacterial density of 10^8.5^ CFU.g^-1^ FW. Fluxes are 0 when stem cannot enrich xylem sap. gS: grams of stem dry weight. D-Glc: D-Glucose, L-Pro: L-Proline, Sucr: sucrose, L-Asp: L-Aspartate, L-Phe: L-Phenylalanine, L-Leu: L-Leucine, L-Val: L-Valine, L-Ile: L-Isoleucine, L-Tyr: L-Tyrosine, L-Thr: L-Threonine.

This enhanced capacity for growth at higher bacterial densities was due to the enrichment of xylem sap in glucose, sucrose, and amino acids, facilitated by the stem’s contribution (Figure 3D; Supplementary Figure 1). Nutrients from the stem thus partially counterbalanced the effect of reduced transpiration and allowed *R. pseudosolanacearum* to reach higher densities. However, the plausibility of such extensive rewiring of stem fluxes by the pathogen is questionable in light of experimental data. Indeed, bacterial population dynamics measured *in planta* (Gerlin et al., 2021) show near-complete cessation of bacterial growth at approximately 10^9^ CFU.g^-1^ FW, aligning more closely with scenario #4 or #5 (without stem resource hijacking) than with scenario #6.

### The fate of putrescine in the *Ralstonia*-tomato interaction

Putrescine fate during plant – *R. pseudosolanacearum* interaction is frequently questioned, as putrescine is abundantly produced by the bacterium *in vitro* and *in planta* (Gerlin, Escourrou, et al., 2021; Lowe-Power et al., 2018; Peyraud et al., 2016) and contributes to virulence by accelerating wilt symptoms (Lowe-Power et al., 2018). We analyzed predicted fluxes of putrescine in our model including its production by *R. pseudosolanacearum* - calibrated to its experimentally measured excretion rate in a xylem-mimicking medium (Baroukh et al., 2025) - and its assimilation by the plant at low and high bacterial densities (10^6.5^ and 10^8.5^ CFU.g^-1^ FW) (Figure 4A).

**Figure 4:**
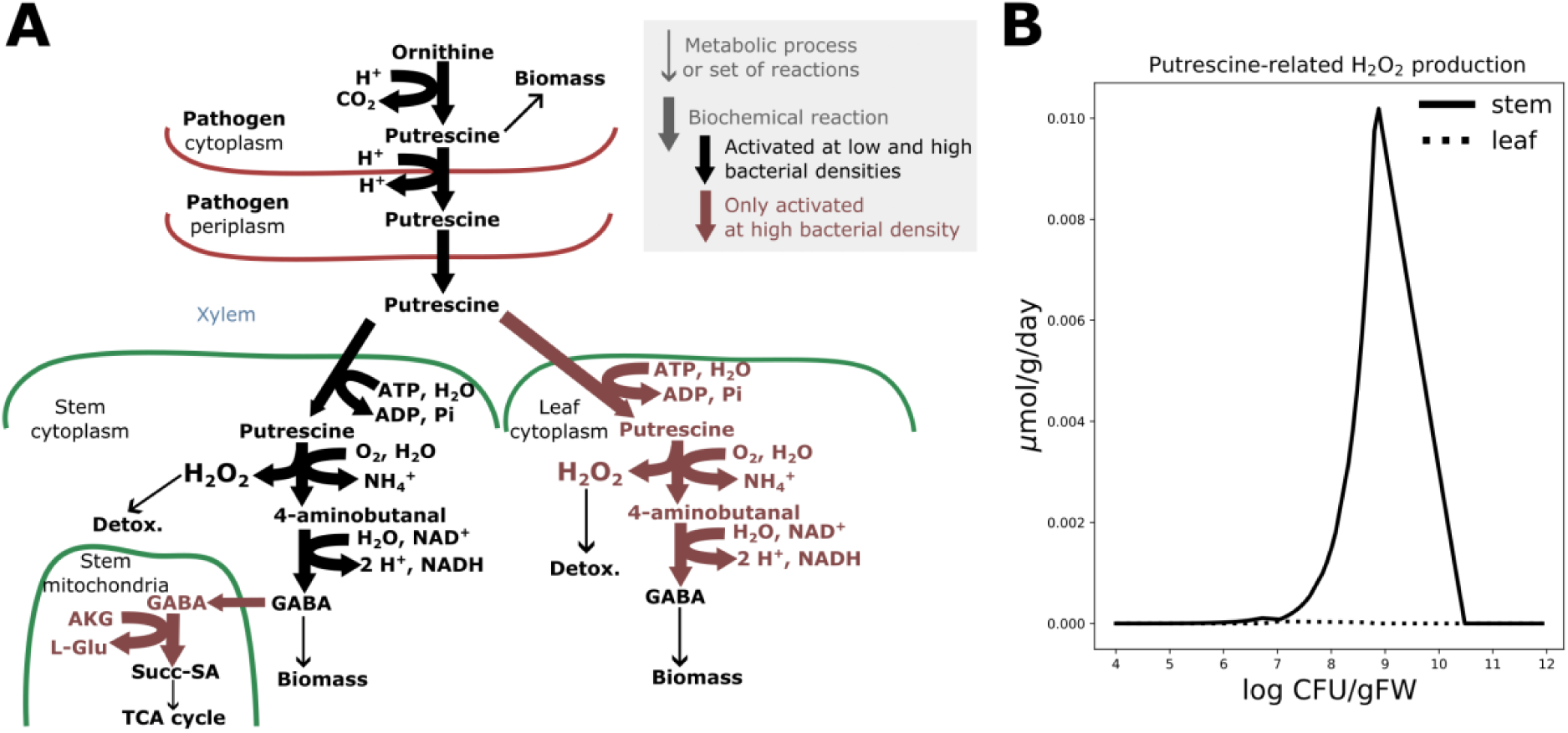
Fluxes predictions surrounding putrescine in VIRION. Only fluxes activated between putrescine production by the pathogen and its assimilation by the plant are represented. In B, g means g of stem (full line) or leaf (dotted line) dry weight.

When putrescine is produced by the bacterium from ornithine, a small portion is incorporated into the bacterial biomass, while the majority is exported to xylem (Baroukh et al., 2022, 2025; Peyraud et al., 2016). Subsequently, the plant aerial tissues can assimilate part of this excreted putrescine. Our model predicted that at low bacterial density, all putrescine is converted into 4-aminobutanal and then into gamma-Aminobutyric acid (GABA). The amount of GABA produced from putrescine at this bacterial density is insufficient to meet the stem biomass requirements for GABA, which are supplemented by other reactions producing GABA from glutamate (glutamate decarboxylase, R_GUDC, EC 4.1.1.15). At high bacterial density, GABA production from putrescine exceeds the stem biomass requirements. This excess is accommodated by: i) activation of the GABA shunt pathway allowing incorporation of GABA into TCA cycle via transport into mitochondria (R_4abut_tx_m_) and conversion into succinate-semialdehyde (4-aminobutyrate transaminase, R_ABTArm) in the stem, ii) activation of the pathway enabling conversion of putrescine into GABA and its assimilation into the leaf. Other reactions associated with putrescine degradation such as conversion into N-acetylspermidine (N1-acetylspermidine oxidase, identifier R_RE1537C in the model) or into N-Methylputrescine (Putrescine N-methyltransferase, R_PMT), were not predicted to carry fluxes, because these reactions produce dead-end metabolites not connected to central metabolism. Interestingly, excess putrescine assimilated by the plant via GABA leads to mandatory production of hydrogen peroxide (H_2_O_2_) (Figure 4B), a key modulator of redox homeostasis (Gerlin, Baroukh, et al., 2021), particularly in the stem. H_2_O_2_ production follows pathogen growth, peaking at approximately at 10^9^ CFU.g^-1^ FW. At higher bacterial densities, the pathogen’s limited growth capacity reduces putrescine production flux, thereby decreasing H_2_O_2_ flux.

## Discussion

### Modeling a plant-pathogen interaction using Flux Balance Analysis

FBA is a methodology widely used in the systems biology community because of its simplicity to set up, its capacity to model at the genome-scale level and its relatively low number of parameters to calibrate (Orth et al., 2010). However, this methodology relies on the quasi-steady-state assumption (QSSA), assuming that no internal metabolites of the system accumulate or get depleted, as well as the fact that the system is optimal with respect to an objective (most often assumed to be growth). The difficulty in modeling a plant-pathogen system lies in the fact that the pathogen grows at the expense of the plant, implying antagonistic objectives between the organisms composing the system. Therefore, questions arise regarding the choice of the objective function that should be optimized. In addition, the pathogen might deplete or accumulate some metabolites upon its growth in the plant, which probably challenges the QSSA.

As a first approximation to quantitatively model a plant-pathogen interaction, we proposed in this work a methodological framework consisting of solving, for several bacterial densities, 4 sequential FBA models in which each FBA model is constrained by the optimal value of the previously solved FBA. In this methodology, we thus assumed that for each bacterial density inside the plant, the internal metabolites were “at steady-state”. We also assumed that pathogen growth was at the expense of the aerial parts of the plant. Despite these strong hypotheses, VIRION was able to illustrate its relevance by predicting growth rates of the aerial part of the plant and the pathogen close to what was experimentally observed. VIRION also allowed the dissection of novel and previously unquantified aspects of the *R. pseudosolanacearum* – tomato interaction, including the critical of transpiration decline in plant growth arrest, the relative importance of carbon compared to other elements, and the potential contribution of stem resource hijacking.

As future directions, the model could be used to study the impact of plant environmental conditions on the plant-pathogen interaction or extended to consider plant defenses via the synthesis of anti-microbial compounds and the resulting bacterial mortality (Zrimec et al., 2025). From a methodological point of view, the QSSA could be slightly relaxed on the metabolites composing the xylem sap, which would allow for more dynamic and relevant modeling of the plant-pathogen interaction.

### Using mathematical modeling to better understand bacterial wilt disease

Since 2016, several mathematical models have been developed to better understand the biology behind bacterial wilt disease. The first model, released in 2016, consisted of a high-quality metabolic network of *R. pseudosolanacearum* simulated in diverse *in vitro* environments using FBA (Peyraud et al., 2016). This model allowed measuring the ATP- equivalent cost of EPS synthesis and virulence. This metabolic network was subsequently interfaced with a regulatory network controlling multiple virulence factors and simulated across a broad set of environmental conditions using a discrete logical modeling method (Peyraud et al., 2018). The resulting model revealed that the virulence regulatory network exerts a control of the primary metabolism to promote robustness upon infection.

More recently, Baroukh et al. (2024) modeled the *in planta* population dynamics of *R. pseudosolanacearum* during tomato infection using a macroscopic modeling framework relying on Monod kinetics. This mathematical model predicted substrate consumption, putrescine production and biomass growth of *R. pseudosolanacearum*.

Unlike previous models, it did not incorporate the metabolic network of *R. pseudosolanacearum* or its regulatory network, nor did it include the plant’s metabolic network, which was simply modeled as a continuous environment providing substrate fluxes to the pathogen. Despite these simplifications, the model accurately reproduced experimental observations, including high bacterial densities and the depletion of glutamine and asparagine. Additionally, it estimated key parameters such as the bacterial mortality rate within the plant and the rate at which bacterial putrescine is assimilated by the plant.

In the present study, the metabolic networks of *R. pseudosolanacearum* and the tomato plant were merged, and the resulting model was simulated using sequential FBAs for diverse bacterial densities within the plant. The model demonstrated several key findings: i) the plant’s photosynthetic capacity is the most critical factor for bacterial proliferation, ii) a reduction in plant transpiration limits and ultimately stops plant and pathogen growths, iii) hijacking stem resources, potentially through virulence factors, can enhance bacterial growth but is not essential for Ralstonia, and iv) pathogen-excreted putrescine is predicted to be directly reused for plant biomass.

Another metabolic model of a plant-pathogen interaction was recently developed by Rodenburg et al. (2019), representing the *Phytophthora infestans* - tomato leaf pathosystem. This study differs from ours in several key aspects, as it employs a qualitative approach, in contrast to our quantitative methodology. This difference is evident in: i) the *P. infestans* biomass equation, which uses equal stoichiometric coefficients, ii) the absence of mass conversion between plant and pathogen fluxes, iii) the unrestricted uptake of all metabolites from tomato cell cytosol by the pathogen, and iv) the lack of calibration for maximal uptake rates for both *P. infestans* and the plant. Roderburg et al. (2019) also utilized a tomato leaf-centered model, which is substantially less complex than our multi-organ model that incorporates a vascular system, and the impact of *P. infestans* growth on plant metabolism and physiology was not simulated. Indeed, all objective functions in their FBA simulations focused exclusively on pathogen growth, whereas our model considers the interplay between pathogen and plant growth.

In each of the Ralstonia mathematical models developed so far, different modeling approaches have been used, different objects have been modeled, various experimental data have been used for calibration, and distinct biological questions have been addressed. Consequently, each model possesses its own specificity and provides unique insights into the mechanisms at play during bacterial wilt disease. Future directions could involve improving existing models by better representing the plant putrescine degradation pathway. Alternatively, models could be made more complex by considering the spatial aspects of infection and representing the effects of abiotic and biotic conditions of the plant on the pathosystem.

Models incorporating plant immunity and its interaction with *R. pseudosolanacearum* virulence factors could also be developed. The first step would be to better represent all the immune-compounds in the metabolic network of the tomato (Zrimec et al., 2025), since the virulence network of *Ralstonia solanacearum* has already been built. However, extensive metabolomics data for both tomato immune metabolites and Ralstonia virulence metabolites will be necessary for calibration to perform quantitative predictions. The second step would be to add the regulatory layer to such model, to better represent triggering of immunity on the plant side and virulence on the pathogen side. This is much more challenging since it requires the hybridization of discrete-regulatory type models with continuous metabolic models (Liu & Bockmayr, 2020; Thuillier et al., 2022). In this study, we focused on a relatively late stage of interaction when the pathogen begins rapid multiplication in host tissues after successful entry and establishment. While the model does not directly account for immunity and virulence mechanisms, it could still successfully evaluate some virulence-related aspects such as the impact of plant resource diversion by the pathogen or the impact of putrescine excretion on the plant H_2_O_2_ production. VIRION can thus be seen as a starting point for the development of more complex plant-bacteria interaction models that explicitly integrate the interplay between metabolism, immunity and virulence mechanisms.

### Plant transpiration is the main factor leading to growth arrest of both the host and the pathogen wilt disease

VIRION allowed to test and identify which physiological or environmental factors are behind plant and pathogen growth arrests during bacterial wilt disease. First, our simulation results showed that the maximal carbon uptake capacity of the plant (through photosynthesis), not nitrogen or iron, set the maximal plant density of Ralstonia cells achievable (Figure 5). However, we also found that carbon, or any other metabolic bottlenecks such as nitrogen and iron, are insufficient to explain the early arrest of plant growth that was experimentally observed in Gerlin et al. (2021). To explain plant growth arrest, limiting transpiration was necessary. This constraint impacts xylem fluxes and hence access to nutrients for the plant aerial parts and later on the pathogen (Figure 5). Interestingly, at densities where this transpiration-limitation is induced, the plant growth was affected immediately, in agreement with experimental observations, contrary to the bacteria which continued to thrive up to several logs higher. This underlines the pathogen’s perfect metabolic adaptation to this challenging environment.

**Figure 5:**
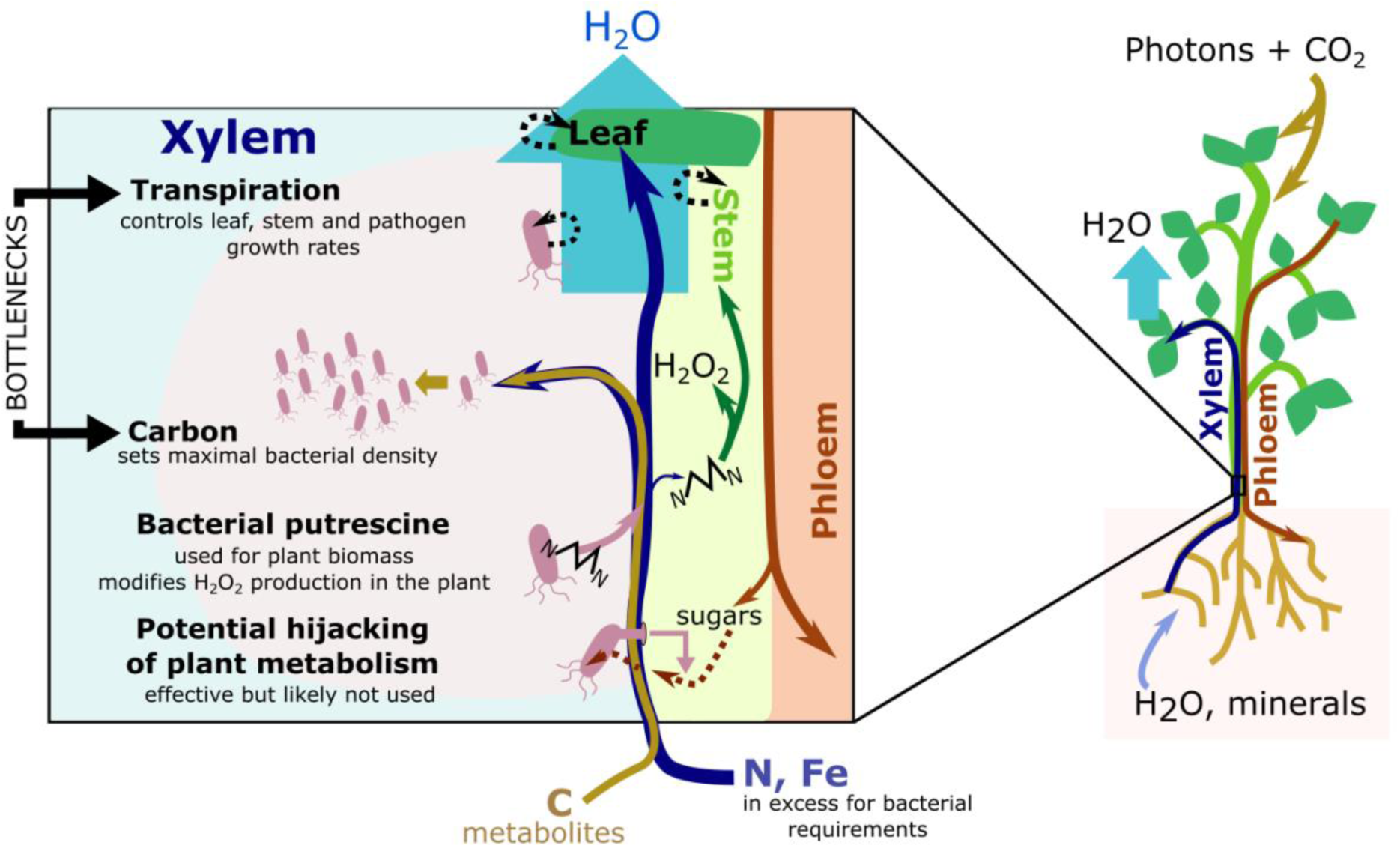
Key metabolic parameters during bacterial wilt at the origin of a successful infection. Decrease in transpiration during infection appears to be the most likely mechanism behind the cessation of plant growth and pathogen growth. Among diverse nutrients, carbon is the most limiting one for bacterial maximal density reachable. The study also showed that, although diverting nutrients from the phloem to the xylem via the stem is an effective strategy for increasing bacterial multiplication in plants, it is poorly used in the case of bacterial wilt. Finally, the effect of bacterial putrescine production on host metabolism was shown to interfere with H_2_O_2_ synthesis in the plant, likely having an effect on ROS response.

Naturally, questions arise on the mechanism responsible of plant transpiration decrease upon *R. pseudosolanacearum* presence in xylem vessels. Physical presence of *Ralstonia* cells could impede xylem flow in itself, but an indirect mechanism such as large bacterial exopolysaccharides excretion could also be responsible by making xylem sap so viscous that it impedes its flow (Ingel, Caldwell, Duong, Parkinson, McCulloh, Iyer-Pascuzzi, McElrone, & Lowe Power, 2022). To discriminate between these hypotheses or others possible mechanisms, further investigation is necessary.

### Stem resource hijacking, a strategy likely used only minimally by *R. pseudosolanacearum* during tomato infection

In the literature, xylem sap is commonly described as a nutrient-poor environment that cannot sustain the biotrophic growth of a plant pathogen to high densities (De La Fuente et al., 2022), thereby requiring the bacterium to force the plant to release additional nutrients, such as sugars (Hamilton et al., 2021; MacIntyre et al., 2020). However, VIRION simulations, integrating both xylem concentrations and flow, tend to show that, even with a transpiration limit, a moderate growth rate of *R. solanacearum* can be reached up to 10^9^ CFU.g^-1^ FW solely through biotrophic growth, in agreement with experimental observed values (Gerlin, Escourrou, et al., 2021). From 10^10^ CFU.g^-1^ FW, a growth rate is still present but very close to 0. Thus, xylem sap, when considered as the product of a continuous flow, is not poor and very high bacterial densities can be reached solely through biotrophic growth of the pathogen.

We then investigated how resource hijacking could contribute to *R. pseudosolanacearum* growth. We confirmed by simulation that this mechanism could theoretically be an effective pathogen strategy to boost its growth and compensate, for example, the effect of transpiration decline, since it allowed the pathogen to have a high growth rate up to 10^10^ CFU.g^-1^ FW. We also confirmed that in the case of stem resource hijacking, this would primarily involve sugars and some amino acids (Figure 5). This would explain why Hamilton et al. (2021) reported a slight increase of sugars in infected xylem sap. Nevertheless, we also showed that this mechanism is probably dispensable for *R. pseudosolanacearum* since pathogen growth almost stops at around 10^9^ CFU.g^-1^ FW and not 10^10^ CFU.g^-1^ FW (see bacterial population dynamics *in planta* in Gerlin et al. (2021)). This also aligns with the observation that mutants defective in sugar assimilation still fully wilt tomato plants, with a slight delay (Hamilton et al., 2021; Gerlin, Escourrou, et al., 2021). We thus conclude that hijacking plant resources from the stem is not likely to occur in bacterial wilt or, if it does, the quantities of nutrients hijacked are much lower than the maximal quantity that can be theoretically hijacked, as predicted by the model.

Reducing the amount of glutamine in the xylem sap, which is the major carbon source for the bacterium, seems to be a better strategy for limiting the growth of *R. pseudosolanacearum* in the plant. However, such modifications would likely affect plant metabolism and phenotype even before impacting the pathogen, as our model tends to indicate. Nevertheless, it might be worthwhile to consider how plant variety of managing the plant’s environment, such as nutrition or watering, could mitigate xylem sap flow and composition and thus minimize *Ralstonia* growth in the plant.

While *R. pseudosolanacearum* appears to favor a fast consumption of resources, as its metabolic and growth capacities outcompete those of the plant, other plant pathogen might not exhibit this behavior but instead favor a reduced growth in xylem vessels. In particular, *Xylella fastidiosa*, although capable of metabolizing various xylem sap components (Gerlin et al., 2020), has very limited growth rates *in vitro* and *in planta*, and is described as adopting a self-limiting behavior (Gerlin et al., 2020; Roper et al., 2019). In these conditions, the plant would be able to dominate the trophic interaction and limit bacterial spread. This suggests that these pathogens would favor a long stay in xylem vessels at low bacterial densities. In this case, a coexistence with the plant takes place in which the bacterium is not competing for resources, and both the plant and the pathogen can remain metabolically active.

## Methods

### Construction of VIRION

The cell tomato metabolic model from Gerlin et al., (2022) was used as a starting point. Reactions for iron, manganese, cobalt, molybdenum and zinc assimilation, which were not represented in the initial metabolic model, were added as these metabolites were necessary for predicting *R. pseudosolanacearum* growth using its genome-scale metabolic model (Peyraud et al., 2016). This addition, as well as other changes or improvements on putrescine metabolism and biomass are fully described in Supplementary Text 1.

From the single-cell tomato model, a multi-organ model was generated using the same procedure as in Gerlin et al., (2022). Briefly, using an in-house script, the metabolic network was replicated into three metabolic networks representing leaf, stem and root with adequate biomass reaction for each organ. Exchange reactions were added to allow the exchange of metabolites between organs through xylem and phloem, considering weight ratios between the different organs. Xylem represented exchanges between root and aerial parts, while phloem represented exchanges between aerial parts and roots. Stem could exchange in both directions with the phloem. Contrary to our previous multi-organ metabolic model, stem could uptake elements from the xylem but could not provide elements to this compartment, except when mentioned. As in Gerlin et al., (2022), a transport cost was added for each exchange reactions except for water. Different physiological constraints (uptakes of water and minerals by roots only, no photosynthesis for root, limited photosynthesis for stem) were integrated to model the different roles of each organ as in Gerlin et al., (2022).

The in-house scripts also merged *R. pseudosolanacearum* metabolic network taken from Peyraud et al. (2016) to the multi-organ network, generating a four-compartment model: leaf, stem, root, pathogen. The boundary metabolites were removed from the metabolic model of *R. pseudosolanacearum*, and external metabolites of the pathogen model were considered as xylem metabolites. To avoid dead-end metabolites, sink reactions were added allowing the exit of *R. pseudosolanacearum* metabolites excreted into the xylem and not present in healthy plant xylem sap. Simulations revealed very few sink reactions were used, and at very low fluxes. In addition, experimental data showed that few metabolites were enriched upon Ralstonia infection in xylem sap (Gerlin, Escourrou, et al., 2021; Lowe-Power et al., 2018). The weight ratio between *R. pseudosolanacearum* and the plant was implemented in the exchange reactions between the xylem and the bacterial periplasm and were modified to represent the different bacterial densities. Conversion of *R. pseudosolanacearum* weight into colony forming unit (CFU) per g of fresh weight (FW) was estimated to follow 1 CFU.g^-1^ stem FW = 1.10 · 10^-11^ g bact / 1.52 g stem dry weight, based on: i) OD/CFU and OD/g ratios from Peyraud et al. (2016), ii) fresh weigh/dry weight (FW/DW) from Gerlin et al. (2022), iii) the leaf/stem and stem/root ratio in tomato metabolic model (Gerlin et al., 2022) which impose relative contents of 1.52 g stem DW and 3.37 g leaf DW for 1 g root DW (Supplementary Text 2).

### Calibration of *R. pseudosolanacearum* metabolic model for growth in planta

To determine growth, excretion and assimilation capacities of *R. pseudosolanacearum* in the presence of xylem carbon sources, we used the data and model generated in Baroukh et al. (2024). In the study, a shake-flask growth experiment was performed in a minimal medium supplemented with a mixture of organic substrates whose relative concentrations mimics a tomato xylem. Growth, substrate assimilation and product excretion were monitored and used to determine uptake, excretion and growth rates among this combination of carbon sources. Glutamine and glucose assimilation followed Monod-type kinetics and were driving growth, while other substrate assimilation rates were proportional to the assimilation of glutamine and glucose. From the Monod kinetics, two maximal assimilation rates, one for glutamine and one for glucose, were implemented as a constraint bound in the model:

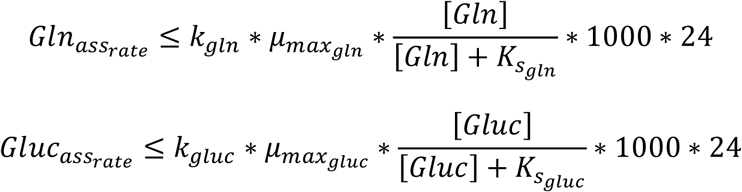

With:

- *Gln*_*ass*rate_ the assimilation rate of glutamine in the model (mmol·gB^-1^·day^-1^), *k*_*gln*_ (mmol·mgB^-1^) the quantity of glutamine necessary to yield 1mg of biomass, *μ*max_*gln*_ (h^-1^) the maximal growth rate on glutamine, *K*_*sgln*_ (mM) the semi-saturation growth term on glutamine, [*Gln*] (mM) the concentration of glutamine in tomato xylem sap as measured in Gerlin et al. (2021)
- *Gluc*_*assrate*_ the assimilation rate of glucose in the model (mmol·gB^-1^·day^-1^), *k*_gluc_ (mmol·mgB^-1^) the quantity of glucose necessary to yield 1g of biomass, *μ*max_gluc_ (h^-1^) the maximal growth rate on glucose, *K*_*s*_ (mM) the semi-saturation growth term on glucose, [*Gluc*] (mM) the concentration of glucose in tomato xylem sap as measured in Gerlin et al. (2021)

From all the other substrates, to ensure assimilation only when glucose or glutamine were assimilated, a constraint proportional to glucose and glutamine rate was added:

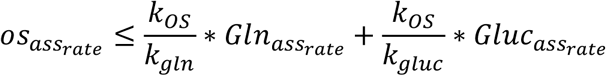

With *OS*_*assrate*_(mmol·gB ·day) the assimilation rate of the other substrate (phenylalanine, valine, leucine, arginine, sucrose, tyrosine, proline, lysine, threonine, isoleucine, asparagine). All values were taken from Baroukh et al. (2024).

The growth-associated and non-growth associated ATP maintenance terms were calibrated so that *R. pseudosolanacearum* growth rate prediction was in agreement with experimental data measured in Baroukh et al. (2024).

### Sequential FBAs

Modeling of the pathosystem was performed using four successive FBAs with different objective functions.

1. Minimization of photon uptake

This first FBA was performed on the plant alone. It minimized photon uptake while leaf, stem and root growth rates were constrained to their experimental values taken from Gerlin et al. (2022). Minimizing photon uptake allowed to estimate the physiological capacities (photosynthesis, iron, nitrogen uptake) of a healthy plant.

1. Maximization of the pathogen growth

The second FBA was performed on the plant-pathogen model. It maximized the pathogen biomass growth while the photon uptake rate was constrained to the value found in FBA 1 (representing healthy plant photosynthetic capacities). This constraint avoided to have an infected plant with a higher photosynthesis rate than a healthy plant. The leaf and stem growth rates were relaxed, to model the fact that *R. pseudosolanacearum* resource acquisition and proliferation can hinder the growth of the aerial part, as observed experimentally (Gerlin, Escourrou, et al., 2021). Root growth rate remained constrained as it was found experimentally that root growth was not impacted by *R. pseudosolanacearum* presence (Gerlin, Escourrou, et al., 2021).

In this FBA, nitrogen or iron uptake rate could also be constrained to the ones of a healthy plant as found in FBA 1. Transpiration rate of the plant could also be constrained. Indeed, measurements of transpiration during tomato – *R. pseudosolanacearum* interaction revealed a decrease of transpiration when the pathogen is present at a density higher than 10^7^ CFU per g of stem fresh weight (FW) (Gerlin, Escourrou, et al., 2021). We estimated thanks to a linear regression the following transpiration decrease linear rule:

reduction_flow_coeff *= -0.2746* [log_10_ CFU.g^-1^ FW] + 2.8764*.

As xylem fluxes of matter depend on the transpiration rate, the upper bound of each flux exchanged between the root and the xylem (v_Exch_r_xyl_i_) was constrained by the value found in FBA1 (vFBA1_Exch_r_xyl_i_) reduced by the reduction_flow_coeff, when the bacterial density exceeded 10^7^ CFU.g^-1^ FW:

If *[log_10_ CFU.g^-1^ FW]* > 7 then v_Exch_r_xyl_i <=_ vFBA1_Exch_r_xyl_i *_ reduction_flow_coeff

1. Maximization of the growth of the aerial parts

The third FBA was performed on the plant-pathogen model. It maximized biomass growth of the stem and the leaves of the plant, while the pathogen growth rate was constrained to the value found in FBA 2. A constant ratio between leaf and stem biomass was imposed to ensure balanced growth of both the stem and the leaves. This FBA determined the growth of the aerial parts achievable in the presence of *R. pseudosolanacearum*, since the pathogen used part of xylem nutrients.

1. Minimization of the sum of absolute fluxes

The fourth and final FBA was performed on the plant-pathogen model. It minimized the sum of all fluxes of the model to ensure a parsimonious enzyme usage in both organisms as well as avoid the presence of Stoichiometric Balanced Cycles. The leaf and stem growth rates were constrained to the values found in FBA 3.

### Software requirements

Simulations were performed with Python 3 scripts, the open access libraries lxml, pandas and the linear programming solver CPLEX Python API, developed by IBM. The scripts used to build the genome-scale metabolic models and perform all the simulations can be found on Github at this link: https://github.com/cbaroukh/VIRION.

An academic or commercial license of CPLEX Python API (ILOG CPLEX Optimization Studio) is mandatory to run the scripts. The academic license is free for academic institutions. The basic “free Community Edition” version of CPLEX is not suitable and will provide an error message.

## Data availability

All data supporting the findings of this study are available within the paper and within its supplementary materials. The scripts used to perform all the simulations, as well as the genome-scale metabolic models, can be found on Github at this link: https://github.com/cbaroukh/VIRION.

## Author contributions

Caroline Baroukh: Conceptualization (lead); Data curation (equal); Formal analysis (equal); Funding acquisition (lead); Investigation (equal); Methodology (equal); Project administration (lead); Resources (equal); Supervision (lead); Writing – original draft (supporting); Writing – review and editing (equal).

Léo Gerlin: Data curation (equal); Formal analysis (equal); Methodology (equal); Investigation (equal); Writing – original draft (lead); Writing – review and editing (equal).

Stéphane Genin: Conceptualization (supporting); Methodology (supporting); Resources (equal); Writing – review and editing (equal).

## Competing interest statement

The authors declare no competing interests.

## Funding statement

Léo Gerlin was funded by a PhD grant from the French Ministry of National Education and Research. The study was funded by the French Laboratory of Excellence (LABEX) project TULIP (ANR-10-LABX-41 and ANR11-IDEX-0002-02). The funders had no role in study design, data collection and analysis, decision to publish, or preparation of the manuscript.

## Supplementary data

**Supplementary Figure 1**

*R. pseudosolanacearum* uptake rates depending on the bacterial density and the limiting factors. g: g of *R. pseudosolanacearum* dry weight. CFU: colony forming units.

**Supplementary Text 1**

Modifications of tomato metabolic model to build a whole plant – pathogen model.

**Supplementary Text 2**

Integration of bacterial density in VIRION.

---

**Supplementary figure 1:**
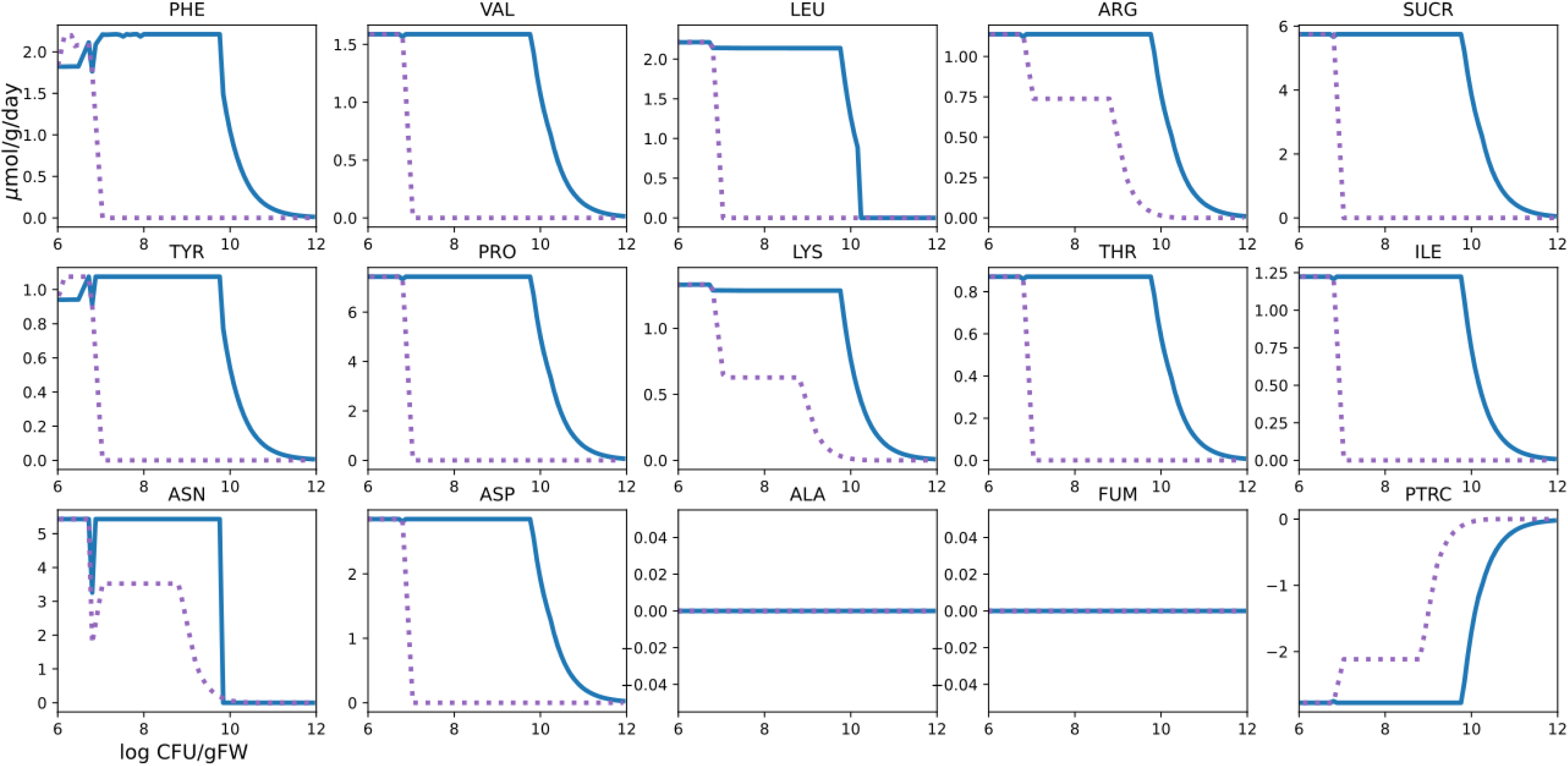
*R. pseudosolanacearum* uptake rates depending on the bacterial density and the limiting factors. g: g of *R. pseudosolanacearum* dry weight. CFU: colony forming units.

**Supplementary Table 1.**
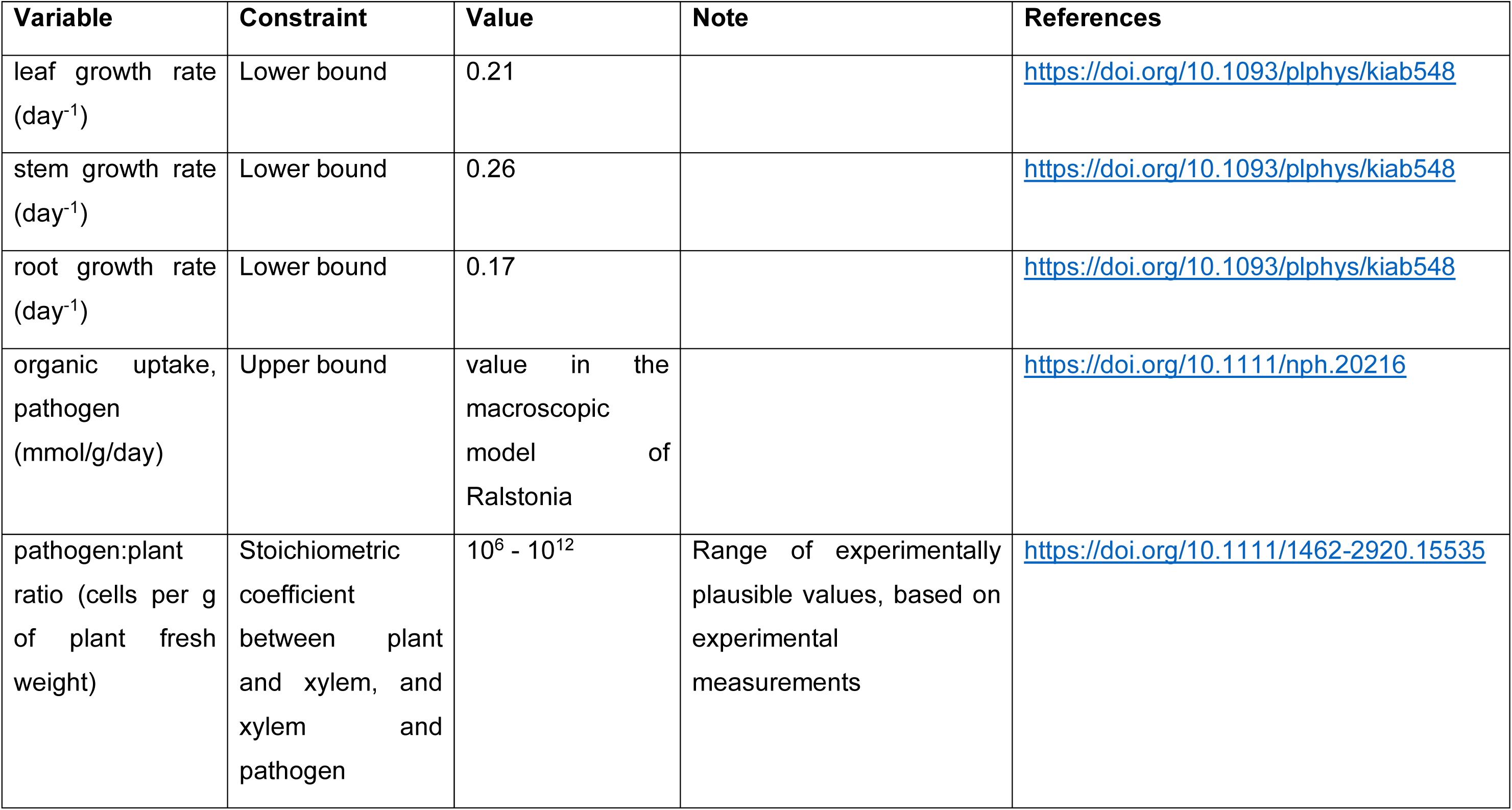
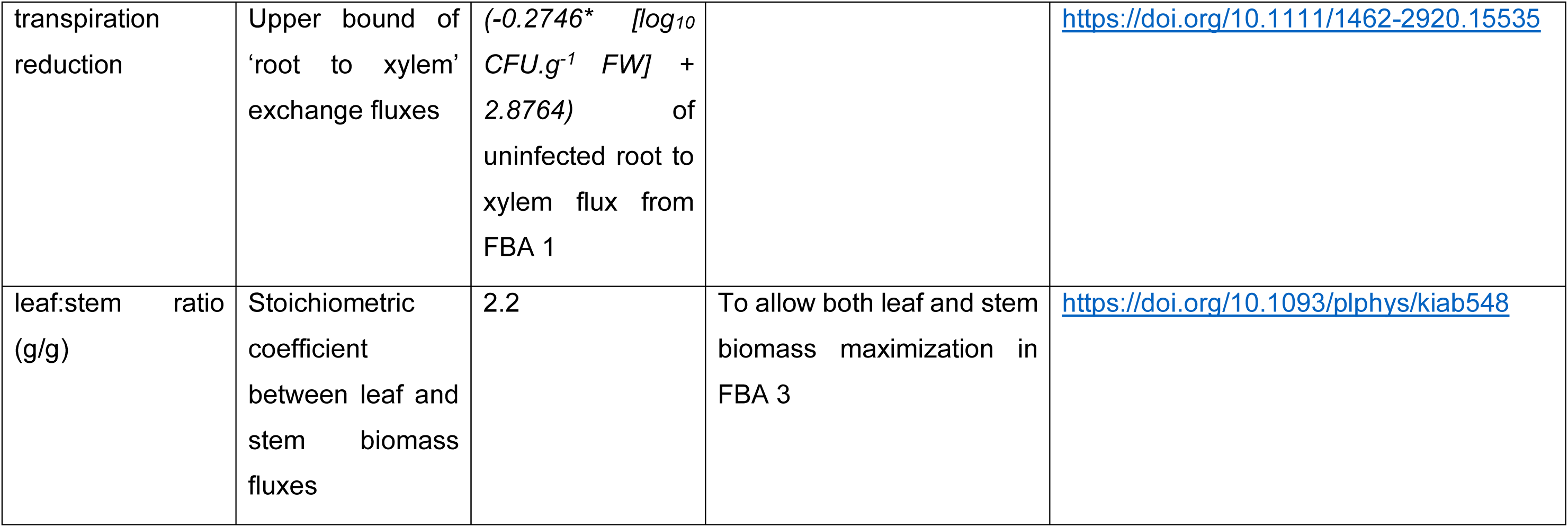
Experimental constraints used to calibrate the multi-organism metabolic model.

## Supplementary text 1

### Modifications of tomato metabolic model to build a whole plant – ***pathogen model***

**1. Determination of iron concentration in tomato plant from literature for integration in biomass**

Iron was initially absent from tomato biomass. Regarding its importance in plant – *Ralstonia* interaction (ref), we added it to organ biomass reactions (R_BIOMASS_ROOT, R_BIOMASS_STEM, R_BIOMASS_LEAF). We determined the coefficient value in the equation following the procedure below.

**· From Twyman, 1959 [1]**

Mean concentration of iron in dry leaf tissue of tomato plants:

[Fe] = 144 ppm at the highest iron supply (Table 2)

[Fe] = 144 ppm = 144 mg Fe/kg leaf DW

Note: at the lowest iron supply, the value remains close (112 mg Fe/kg leaf DW).

**· From Carpena et al., 1988 [2]**

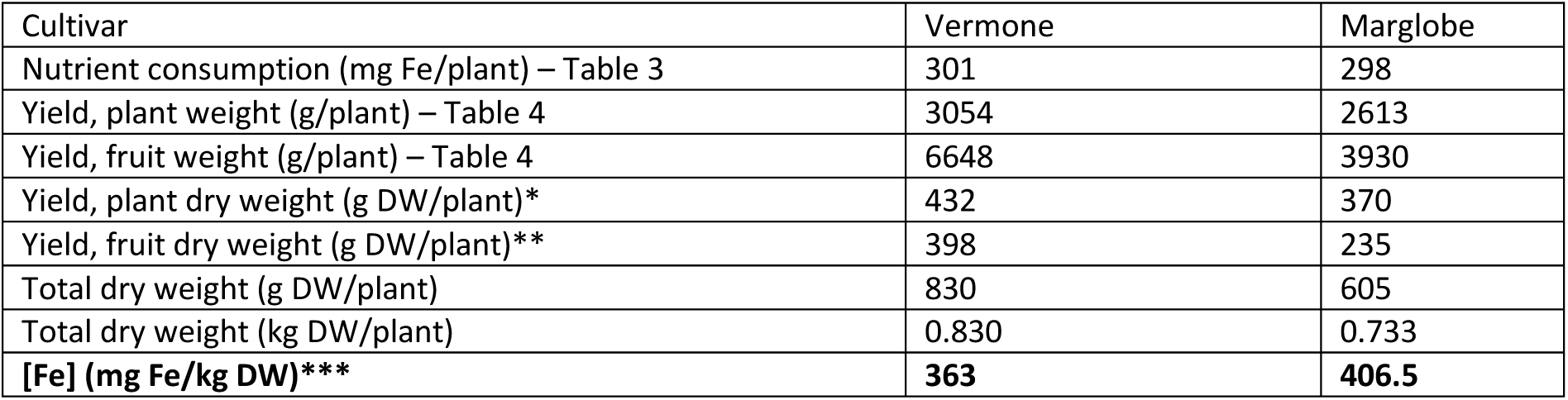

→ **Integration of iron concentration in tomato model**

The range estimated from literature is **144 – 406.5 mg Fe/kg DW** in a tomato plant. Biomass equation represents 1 g of organ DW, corresponding to 1.44 – 4.07 10^-4^ g Fe. Using the atomic weight of Fe (55.845 g/mol), it represents 2.58 – 7.28 10^-6^ mol Fe.

Biomass equation is expressed in µmol/gDW, so the range of values to integrate in the model is: **2.58 – 7.28 µmol/gDW.** Iron is majorly found in and between plant organs as Fe2+ (M_fe2_c) [5], so iron will be incorporated in biomass equation as M_fe2_c (see reference below).

Here, we do not take into account the organ specificity of iron concentration and add the same value in leaf, stem and root. We integrate the highest value to mimic a maximal requirement. Simulations presented in the paper show that even this high value has poor impact on the interaction.

**2. Addition of minerals required for pathogen growth**

Molybdate (M_mobd_c), manganese (M_mn_c) and cobalt (M_cobalt_c) are present in *R. pseudosolanacearum* biomass equation while they were not modeled in plant initial metabolic model (Sl2183).

We added:

- **For molybdate**

The internal metabolite M_mobd_c was already present. We added the boundary metabolite M_mobd_b and the uptake reaction R_EX_mobd_c, written as follow:

M_mobd_b <-> M_mobd_c

- **For manganese**

We added the internal and boundary metabolites M_mn2_b and M_mn2_c and the uptake reaction R_EX_mn2_c, written as follow:

M_mn2_b <-> M_mn2_c

- **For cobalt**

We added the internal and boundary metabolites M_cobalt2_b and M_cobalt2_c and the uptake reaction R_EX_cobalt2_c, written as follow:

M_ cobalt2_b <-> M_ cobalt2_c

**3. Corrections of amino acid coefficients in biomass equation**

The coefficients for some amino acids were inverted in the previous files of tomato model published in 2022 [3]. While it does not have significant impact on the simulations, either here or in the previous work, we rewrote the coefficients for the following amino acids: M_cys L_c, M_ala B_c, M_his L_c, M_ile L_c, M_leu L_c, M_lys L_c with the correct values from the list of biomass components (Supp File 1 of Gerlin et al. 2022 [3], sheets 3, 4, 5).

**4. Putrescine metabolism**

We manually examined putrescine catabolism in Sl2183, as our initial model was not representing putrescine catabolism. We looked at canonical putrescine catabolism in plants in literature [6] and databases (https://pmn.plantcyc.org/pathway?orgid=ARA&id=PWY-2).

We found that the first reaction of putrescine catabolism, putrescine oxidase* (EC 1.4.3.10) was already included in our model (converting putrescine into 4-aminobutanoate), but we missed the following reaction, aminobutyraldehyde dehydrogenase** (EC 1.2.1.19), converting 4-aminobutanoate into GABA. We found two orthologs (SOLYC06G071290 OR SOLYC03G113800) to the A. thaliana gene (ALDH10A9 / AT3G48170) annotated as aminobutyraldehyde dehydrogenase in BioCyc. This provides evidence for the completeness of the pathway in tomato, and we thus added aminobutyraldehyde dehydrogenase in the modeL

*Putrescine oxidase

R_PTRCOX1 http://bigg.ucsd.edu/universal/reactions/PTRCOX1 in BiGG

PUTRESCINE-OXIDASE-RXN https://biocyc.org/reaction?orgid=META&id=PUTRESCINE-OXIDASE-RXN in BioCyc Reaction in BiGG: M_h2o_c + M_o2_c + M_ptrc_c -> M_4abutn_c + M_h2o2_c + M_nh4_c

**Aminobutyraldehyde dehydrogenase

R_ABUTD http://bigg.ucsd.edu/models/iJO1366/reactions/ABUTD in BiGG

AMINOBUTDEHYDROG-RXN https://biocyc.org/reaction?orgid=META&id=AMINOBUTDEHYDROG-RXN in BioCyc Reaction in BiGG: M_4abutn_c + M_h2o_c + M_nad_c -> M_4abut_c + 2.0 M_h_c + M_nadh_c

## Supplementary Text 2

### Integration of bacterial density in VIRION

***Numerical values used***

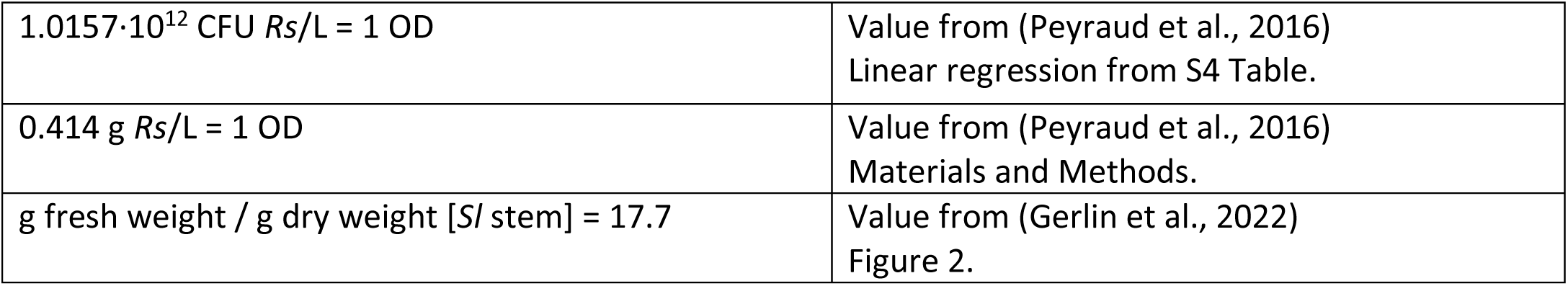

**g / CFU [*Rs*]** = 0.414 g *Rs*/L / 1.0157·10^12^ CFU *Rs*/L = 4.076·10^-13^

**1 cfu/g stem FW** = 1·(g fresh weight / g dry weight [*Sl* stem]) = 17.7 **cfu/g stem DW**

*In VYTOP model, the relative proportions are: 1.52 g stem DW and 3.37 g leaf DW for 1 g root DW. The bacterial density must be determined for 1.52 g stem DW*.

**1 cfu/g stem FW** = 1.52 · 17.7 cfu/1.52 **g stem DW** = 26.904 cfu/1.52 g stem DW

**1 cfu/g stem FW** = 26.904 · (g / CFU [*Rs*]) **g bact / 1.52 g stem DW**

**1 cfu/g stem FW** = 26.904 · 4.076.10^-13^ **g bact / 1.52 g stem DW**

**1 cfu/g stem FW** = 1.10 · 10^-11^ g bact / **1.52 g stem DW**

*Rs*: *Ralstonia pseudosolanacearum* GMI1000

*Sl*: *Solanum lycopersicum* M80

## Notes

https://github.com/cbaroukh/VIRION

### Competing Interest Statement

The authors have declared no competing interest.

### Summary of Updates

Figures and text revised for clarity

